# Opportunities and limits of combining microbiome and genome data for complex trait prediction

**DOI:** 10.1101/2020.10.05.325977

**Authors:** Miguel Pérez-Enciso, Laura M. Zingaretti, Yuliaxis Ramayo-Caldas, Gustavo de los Campos

## Abstract

The analysis and prediction of complex traits using microbiome data combined with host genomic information is a topic of utmost interest. However, numerous questions remain to be answered: How useful can the microbiome be for complex trait prediction? Are microbiability estimates reliable? Can the underlying biological links between the host’s genome, microbiome, and the phenome be recovered? Here, we address these issues by (i) developing a novel simulation strategy that uses real microbiome and genotype data as input, and (ii) proposing a variance-component approach which, in the spirit of mediation analyses, quantifies the proportion of phenotypic variance explained by genome and microbiome, and dissects it into direct and indirect effects. The proposed simulation approach can mimic a genetic link between the microbiome and SNP data via a permutation procedure that retains the distributional properties of the data. Results suggest that microbiome data could significantly improve phenotype prediction accuracy, irrespective of whether some abundances are under direct genetic control by the host or not. Overall, random-effects linear methods appear robust for variance components estimation, despite the highly leptokurtic distribution of microbiota abundances. Nevertheless, we observed that accuracy depends in part on the number of microorganisms’ taxa influencing the trait of interest. While we conclude that overall genome-microbiome-links can be characterized via variance components, we are less optimistic about the possibility of identifying the causative effects, i.e., individual SNPs affecting abundances; power at this level would require much larger sample sizes than the ones typically available for genome-microbiome-phenome data.

**Author summary:** The microbiome consists of the microorganisms that live in a particular environment, including those in our organism. There is consistent evidence that these communities play an important role in numerous traits of relevance, including disease susceptibility or feed efficiency. Moreover, it has been shown that the microbiome can be relatively stable throughout an individual’s life and that is affected by the host genome. These reasons have prompted numerous studies to determine whether and how the microbiome can be used for prediction of complex phenotypes, either using microbiome alone or in combination with host’s genome data. However, numerous questions remain to be answered such as the reliability of parameter estimates, or which is the underlying relationship between microbiome, genome, and phenotype. The few available empirical studies do not provide a clear answer to these problems. Here we address these issues by developing a novel simulation strategy and we show that, although the microbiome can significantly help in prediction, it will be difficult to retrieve the actual biological basis of interactions between the microbiome and the trait.

## Introduction

The relevance of microbial ecosystems associated with humans and animals in health and production is now widely recognized, e.g., [1–5]. To quantify its influence, the fraction of variance of a given trait explained by the microbiome has been named ‘microbiability’ (*b*^*2*^) [6], in symmetry with the classical ‘heritability’ (*h*^*2*^) concept [7]. Previously, the term “hologenome” had been coined to describe the joint action of genome and microbiome in explaining an observed phenotype [8].

A consequence of microbiability being typically larger than zero is that it can be used to predict complex phenotypes, be it a disease or productive traits. This is an important issue since the use of microbiome data has the potential to alter how medical diagnosis in humans or breeding decisions agricultural species are performed. Several studies have demonstrated the potential value of microbiome data for complex-trait prediction. For example, Rothschild et al. [9] showed that microbiome can be used to improve accuracy in the prediction of obesity and many other phenotypes in humans. Likewise, Lloyd-Price et al. showed that microbiome-data was predicted if future outbursts of bowel disease [10]. In cattle, various studies have shown the predictive power of microbiome for methane emission from rumen microbiome [4,11], feed efficiency and carcass traits in pigs [12,13], and various plant phenotypes (e.g., crop yield and diseases predicted from the microbiota data from the rhizosphere, [14]). On the other hand, since the groundbreaking study of Meuwissen, Hayes and Goddard [15], the prediction of complex traits using genome information has been embraced in both plant [16] and animal breeding [17] as well as in human genetics [18]. Therefore, a natural step further is combining host’s genome and microbiome information to improve complex-trait prediction, a topic that is currently receiving much attention [12,19].

Importantly, microbiome composition can be affected by the host’s genome. For instance, Wang et al. [20] argue that it is evolutionarily justified that the microbiome is under partial host genetic control since a non-negligible fraction of cells in an adult body is made up of microbes, especially in the gut. Beginning with the seminal work by Pomp’s team [21], several studies have confirmed the relationship between host’s genotype and microbiome composition, e.g., [20,22,23]. These microbiome genome-wide association studies (mGWAS) suggest that microbiome abundances can be treated as any other complex trait in humans or livestock [22]. For instance, Crespo-Piazuelo et al. [24] or Ramayo-Caldas *et al*., [25,26] identified several quantitative trait loci (QTL) that modulate gut bacterial and eukaryotic communities. In general, although the ‘heritability’ of each genera or OTU (Operational Taxonomic Unit) is typically weak, considering the whole microbiome simultaneously should increase power [27].

Large scale studies in humans suggest a predominant role of the environment in shaping the gut microbiome [9]. However, regardless of the relative importance of genetic and environmental factors in shaping the microbiota, microbiome composition *per se* can have predictive value. Yet, the use of microbiota for prediction of future phenotypes/disease outcomes, require some level of stability of the microbiome throughout time. In the case of the gastrointestinal tract, microbiota colonization starts at birth, where vertical transmission through the mother’s birth canal occurs. Afterward, microbiota diversity and richness tend to increase as the host ages and reaches stability at adulthood [28,29]. In ruminants, populations inhabiting the rumen progressively appear after birth and partly persists throughout life [30].

As noted, the genome-microbiome-phenome is a complex system; understanding the links between host-genome, microbiota, and phenotypes is an important step towards the effective use of microbiome data for complex trait prediction. In all, despite published reports, we still lack detailed guidelines on the joint usage of microbiome and genome information for complex trait prediction, and on the reliability of parameter inferences. We are ignorant of the number of genes affecting microorganism abundance that can be confidently identified, or on how many microorganism taxa can influence a given phenotype. With this work, we aim to contribute to this important topic focusing on three inter-related questions:

1. How useful can the microbiome be for complex trait prediction?
2. Are microbiability estimates reliable?
3. Can the underlying biological genome-microbiome-links be inferred at a system-level? In a more refined level, can microbiome groups (e.g., OTUs, genera) with sizable causal effects on phenotypes be identified with the typical size of microbiome data sets?

In this study, we address the questions mentioned above via a novel simulation strategy that uses real microbiome and genotype data as input and proposing a variance-component approach which, in the spirit of mediation analyses, quantifies the proportion of phenotypic variance explained by genome and microbiome, and dissects it into direct and indirect effects. Importantly, the approach allows simulating a partial genetic control of host’s genome on the microbiome. This is accomplished using a partial permutation approach that preserves the distribution of the genome and microbiome. We use Bayesian variable selection models to estimate parameters which contemplate the possibility that some or all the features available in the genome and/or the microbiome, have no effects on the trait of interests. We investigate the questions presented above across diverse scenarios regarding the links between host genomes and microbiomes, and of their relations with a complex trait.

## Results and Discussion

The exact nature of the links between genome (**G**), microbiome (**B**), and phenotype (**y**) are largely unknown and will likely vary from case to case. However, we will use the six generic causal models (‘scenarios’) depicted in Fig 1 to shed light on the nature of the genome-microbiome-phenome links. In the ‘Null’ scenario, there is no link between any of the data-layers; while this is unlikely, it serves as an ‘overall null hypothesis’ and it is useful to assess potential biases in parameter estimates. Model ‘Genome’ assumes that **G** only affects the phenotype. In turn, only **B** has a direct effect on phenotype in ‘Microbiome’ and ‘Indirect’ scenarios. The Indirect scenario, however, allows for some of the causative abundances to be controlled genetically. This would be similar to a scenario where a phenotype is directly controlled by gene expression levels and expression in turn is controlled genetically [31,32]. The ‘Joint’ scenario is the simplest configuration for a trait under the influence of both genes and microbiome. It assumes microbiome and genome are independent and that their effects on the phenotype are also independent. The Joint model is the most widely assumed, implicitly, or explicitly, in the literature, e.g., [4,9,12]. The ‘Recursive’ model is similar to the Joint model; however, the Recursive model contemplates the possibility that some causative OTU may be under partial genetic control by the host. Therefore, in this case, the genome has both direct and indirect (microbiome-mediated) effects on phenotypes. Note the Recursive model does not assume that the same loci have simultaneously direct and indirect effects, neither it assumes that all OTU abundances are under genetic control.

**Fig 1.**
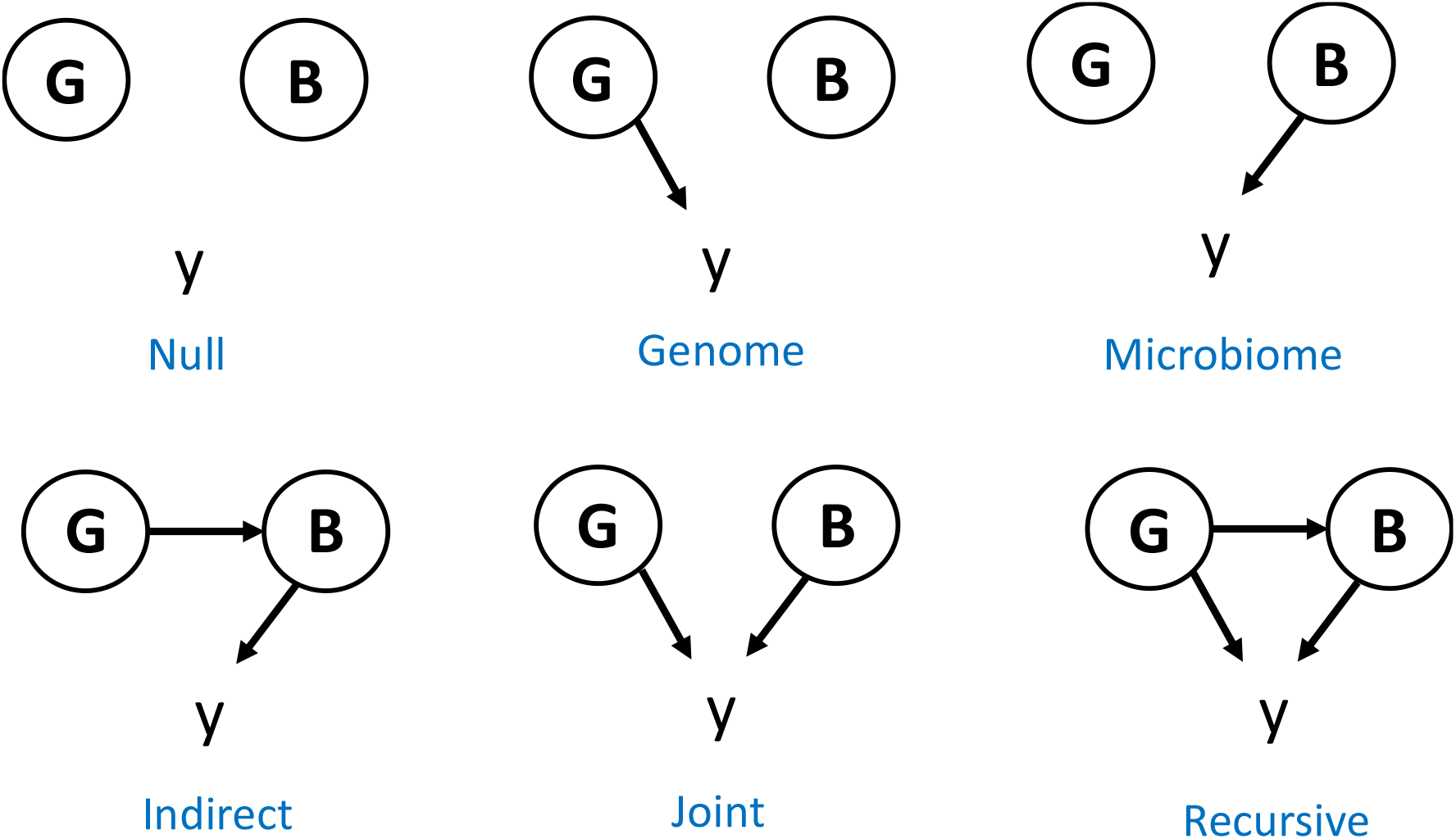
Representation of the scenarios evaluated: **G**, genome, typically comprises marker data; **B**, microbiome; **y**, phenotype of interest; arrows indicate causality. An arrow from **G** to **y** indicates that there is a subset of **G** elements (causative SNPs) that influence **y**; an arrow from **G** to **B** indicates there exists a subset of **G** that influences a subset of abundances in **B** which, in turn, may also influence **y**. An arrow departing from **B** indicates there is a subset of microbial abundances (the causative abundances) that influence **y**. The SNPs affecting **B** need not necessarily be the same SNPs affecting **y** directly in the Recursive scenario. Note **B** can contain one or more sets of abundances such as archaea and bacteria communities, or different time or site sampling points. Without loss in generality, we assume **B** is a single community.

We use the causal models depicted in Fig 1 to simulate genome-microbiome-phenotype data using different configurations regarding the number of causative loci (QTN) and the number of OTUs with effects on phenotypes, as well as the number of OTUs that were affected by host’s genome. Table 1 summarizes the simulation models and parameter values.

**Table 1.** Definition of scenarios evaluated and parameters chosen: **G**, genome; **B**, microbiome; **y**, phenotype of interest; N_QTN_, number of SNPs with a direct causal effect on **y**; N_OTU_, number of OTUs with a direct effect on **y**; N_OTU(g)_, number of OTUs with a direct effect on **y** that are genetically determined, i.e., they are a subset of N_OTU_; *h*^*2*^ is heritability, *b*^*2*^ is microbiability, and *r*^*2*^ = *h*^*2*^ + *b*^*2*^.

| Scenario | Abbreviation | $N_{\text{QTN}}$ | $N_{\text{OTU}}$ | $N_{\text{OTU(g)}}$ | $r^2$ | $h^2$ | $b^2$ |
| --- | --- | --- | --- | --- | --- | --- | --- |
| Null | 0 | - | - | - | 0 | 0 | 0 |
| Joint | J | 100 | 25 | 0 | 0.25 | 0.125 | 0.125 |
|  |  |  |  |  | 0.50 | 0.25 | 0.25 |
| Genome | G | 100 | 0 | 0 | 0.25 | 0.25 | 0.00 |
|  |  |  |  |  | 0.50 | 0.50 | 0.00 |
| Microbiome | M | 0 | 25 | 0 | 0.25 | 0.00 | 0.25 |
|  |  |  |  |  | 0.50 | 0.00 | 0.25 |
| Recursive | R | 100 | 25 | 25 | 0.25 | 0.125 | 0.125 |
|  |  |  |  |  | 0.50 | 0.25 | 0.25 |
| Indirect | I | 0 | 25 | 25 | 0.25 | 0.00 | 0.25 |
|  |  |  |  |  | 0.50 | 0.00 | 0.50 |

### A novel data-driven strategy to generate microbiome-genome-phenotype experiments

Two facts make it the simulation of scenarios in Fig 1 challenging: (i) microbiome data follow zero-inflated highly leptokurtic multivariate distributions [33,34], it is not obvious how to sample from these distributions *conditionally* on genome data as required in the Recursive and Indirect scenarios; and (ii) it is difficult to obtain accurate estimates of key parameters, such as microbiability values, in the absence of large scale published – and public – datasets. To circumvent, or at least to alleviate, these constraints we use real data for both **G** and **B**. Specifically, we used publicly available data from two of the largest microbiome studies in livestock, genome data were downloaded from [11] and OTU abundances from [4].

Fig 2 recapitulates the simulation strategy. Full details are given in Material and Methods section, and R code to replicate the analyses are in https://github.com/miguelperezenciso/simubiome). We assume the effects of the causative microbiome abundances are additive on the log scale. Simulation under the Joint scenario is straightforward, since **G** and **B** act independently: sample a list of causative SNPs and abundances, simulate their effects, and apply Eqn. 1 (Material and Methods) to generate phenotype values given observed genotypes and abundances. The case of Recursive and Indirect scenarios is not that obvious because causative abundances are under genetic control and a link must exist between **G** and **B** (Eqn. 2 in Material and Methods). We solved this issue by rearranging abundances within individuals such that the desired correlation between abundance and individual’s genotypes is attained (see Algorithm in Box 1 and R-code in https://github.com/miguelperezenciso/Simubiome/blob/master/sortCor.R). This strategy has the important advantage that the distribution of abundances is not changed.

**Fig 2:**
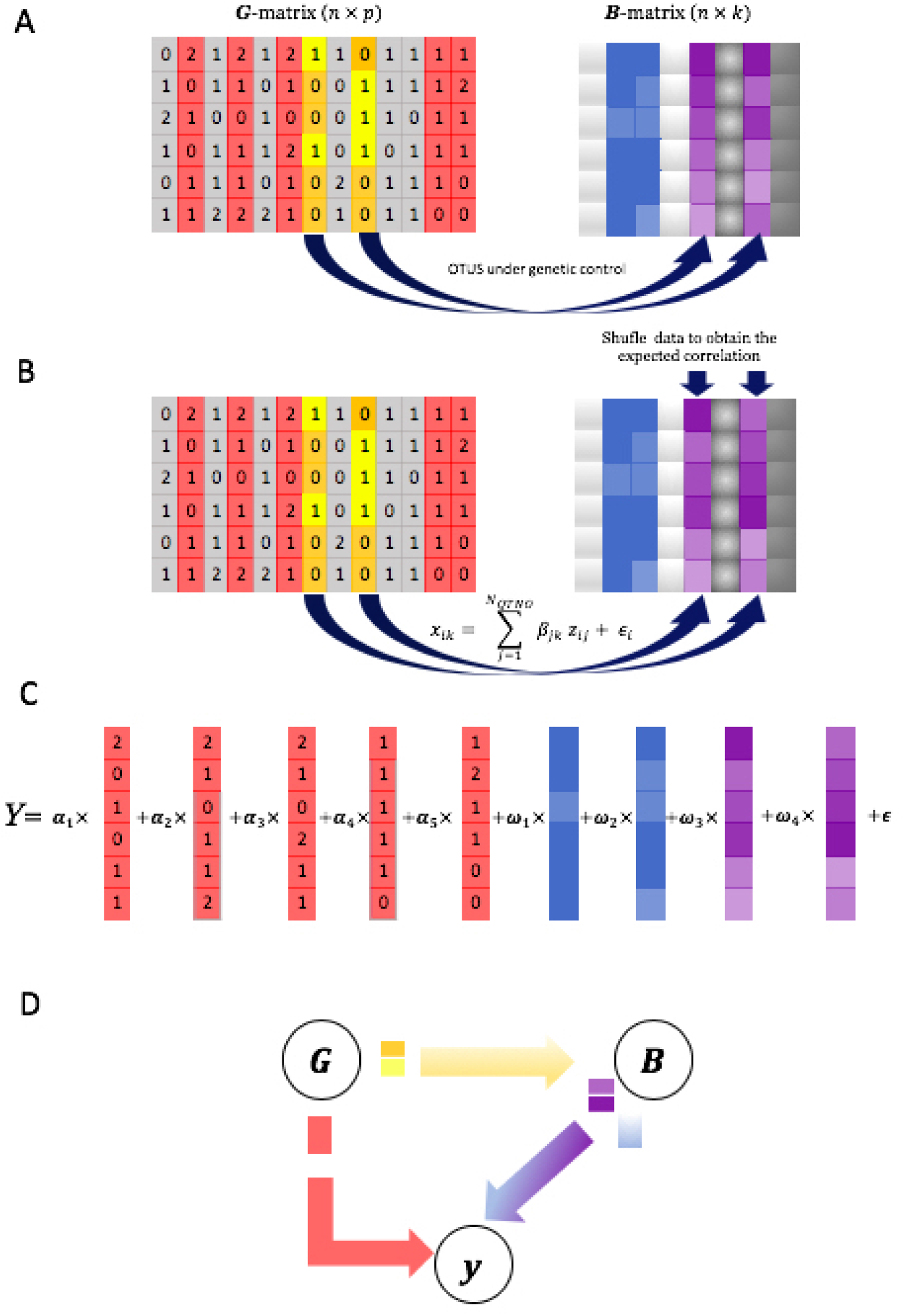
Simulation scheme for the Recursive scenario, i.e., the most complex scenario (Fig 1). **A**) Real input data comprises p genotypes (**G** matrix) and k taxa abundances (**B** matrix). SNPs in grey are neutral, those in red act directly on the phenotype **y**, and those in yellow/orange influence some OTU abundances (marked in magenta color in **B** matrix); abundances in blue are not genetically controlled. **B**) Given simulated effects, a genotypic value controlling the abundances is obtained via Eqn. 2. To fulfill the required heritability, abundances in magenta are reordered; high abundances (represented by a darker color) are associated with genotype ‘1’ just to simplify visualization. A single SNP is shown as causative for each of the two OTUs but there is no limit in practice. **C**) The phenotype is simulated by adding the genome and the microbiome contributions plus a residual. **D)** The general causal diagram is shown.

### How useful can microbiome be for complex trait prediction?

This depends on how much phenotypic variance is jointly explained by the genome (*h*^*2*^) and the microbiome (*b*^*2*^), but also on how efficiently methods capture the relationship between the microbiome and the phenotype, and on how stable the microbiome is. Note prediction accuracy is conditionally independent of whether the microbiome itself is heritable or not. This means that, *given* observed abundances **B** and observed genotypes **G**, it does not matter whether the biological process generating **B** is affected by **G**. In other words, prediction should not be affected by whether the Joint or Recursive scenarios hold, for a constant *r*^*2*^ = *h*^*2*^ + *b*^*2*^. The implications for breeding, however, could be dramatically different. Breeding schemes targeting the microbiome could be designed provided the Recursive scenario holds but make no sense under the Joint scenario.

We compared predictive performance of Bayes C [15] when both genome and microbiome are employed in the model (*Bayes Cgb*) only genome (*Bayes Cg*), or only microbiome data (*Bayes Cb*). First, we verified the null model resulted in no false predictive accuracies (Fig S1A). Fig 3 shows simulated predictive accuracies for the two *r*^2^ values considered (0.25 and 0.50) and for each causative scenario (Fig 1). Predictive accuracies using *Bayes Cgb* were consistently the best. As expected, this was especially the case when both *h*^*2*^ and *b*^*2*^ are larger than zero, that is, when Joint or Recursive scenario hold. In these scenarios, using both sources of variation clearly improved prediction compared to using only genome (*Bayes Cg*) or microbiome data (*Bayes Cb*). Importantly, predictive accuracy was somewhat lower in Joint and Recursive scenarios than in Microbiome or Genome scenarios. This indicates that predictive accuracy does not depend only on total *r*^*2*^, but also on how this variance is split between genome and microbiome. Although this likely occurs because of the larger noise in Recursive or Joint scenarios than in Microbiome or Genome scenarios, it also suggests that our analysis strategy may not be optimum. There is room to develop more efficient tools, especially when the Recursive scenario holds. Note that variance of prediction was larger in the Recursive than in the Joint scenario, i.e., the fact that some abundances are inherited is an additional source of noise.

**Fig 3.**
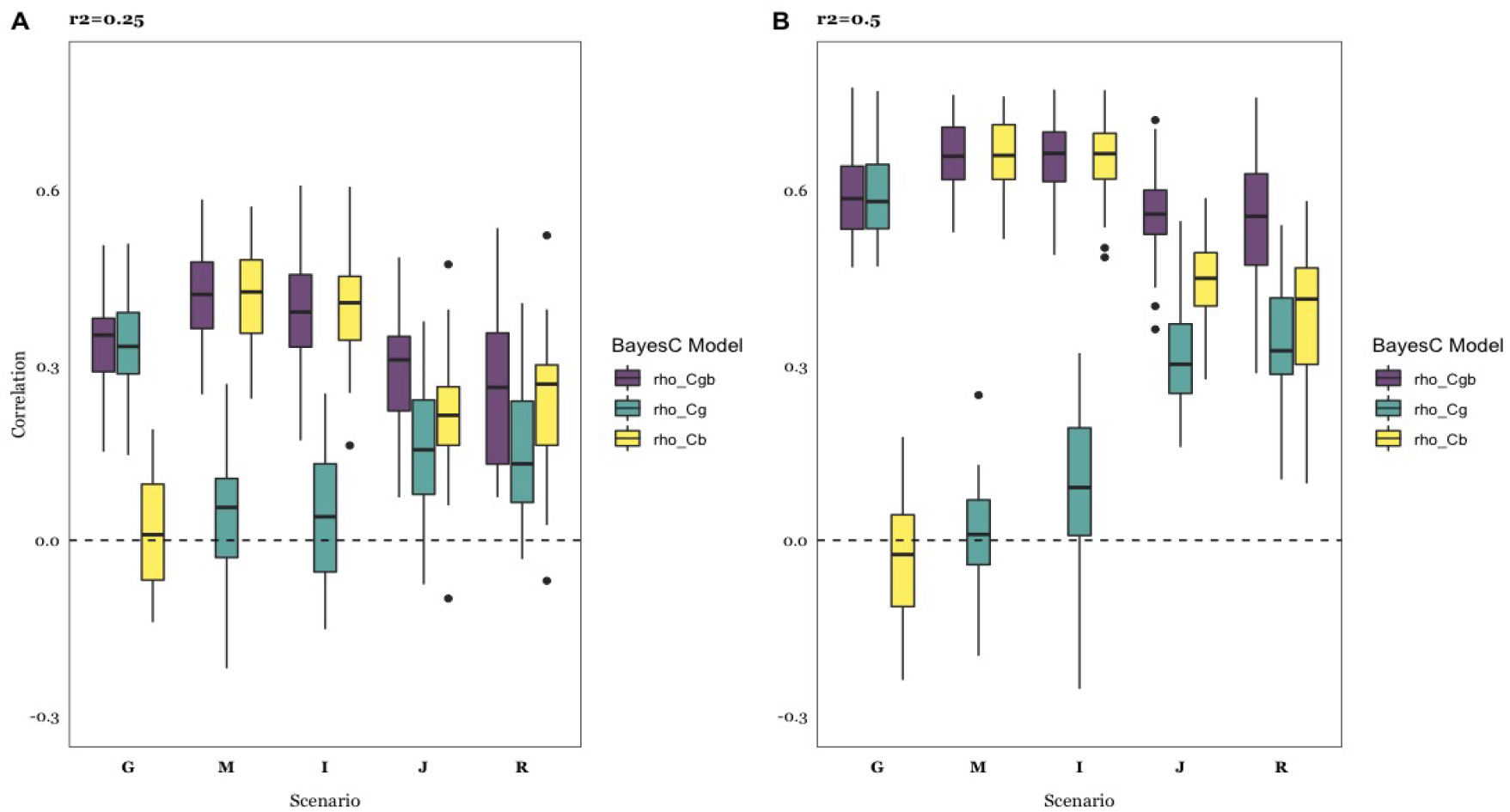
Predictive accuracy, computed as correlation between predicted and observed phenotypes across causal scenarios (Fig 1), for each of the Bayes C models: Cgb considers microbiome and genome; Cg includes genome data only, and Cb includes microbiome data only. **A**: r^2^ = 0.25; **B**: r^2^= 0.50. Details of scenarios are in Table 1: G, Genome; M, Microbiome; I, Indirect; J, Joint; R, Recursive. Results are average of 30 replicates per case.

It is noticeable that predictions were better when only the microbiome influenced the phenotype than when the genome was the only source of variation, a phenomenon also observed with real data [11,12,19]. In this simulation, this occurs likely because the number of causative effects and of input variables (SNPs vs. OTUS) is smaller in the Microbiome or Indirect scenarios than in the Genome scenario. In fact, we do observe a consistent negative correlation between the number of causative OTUs and predictive accuracy in both Joint and Recursive scenarios (Fig 4A).

**Fig 4.**
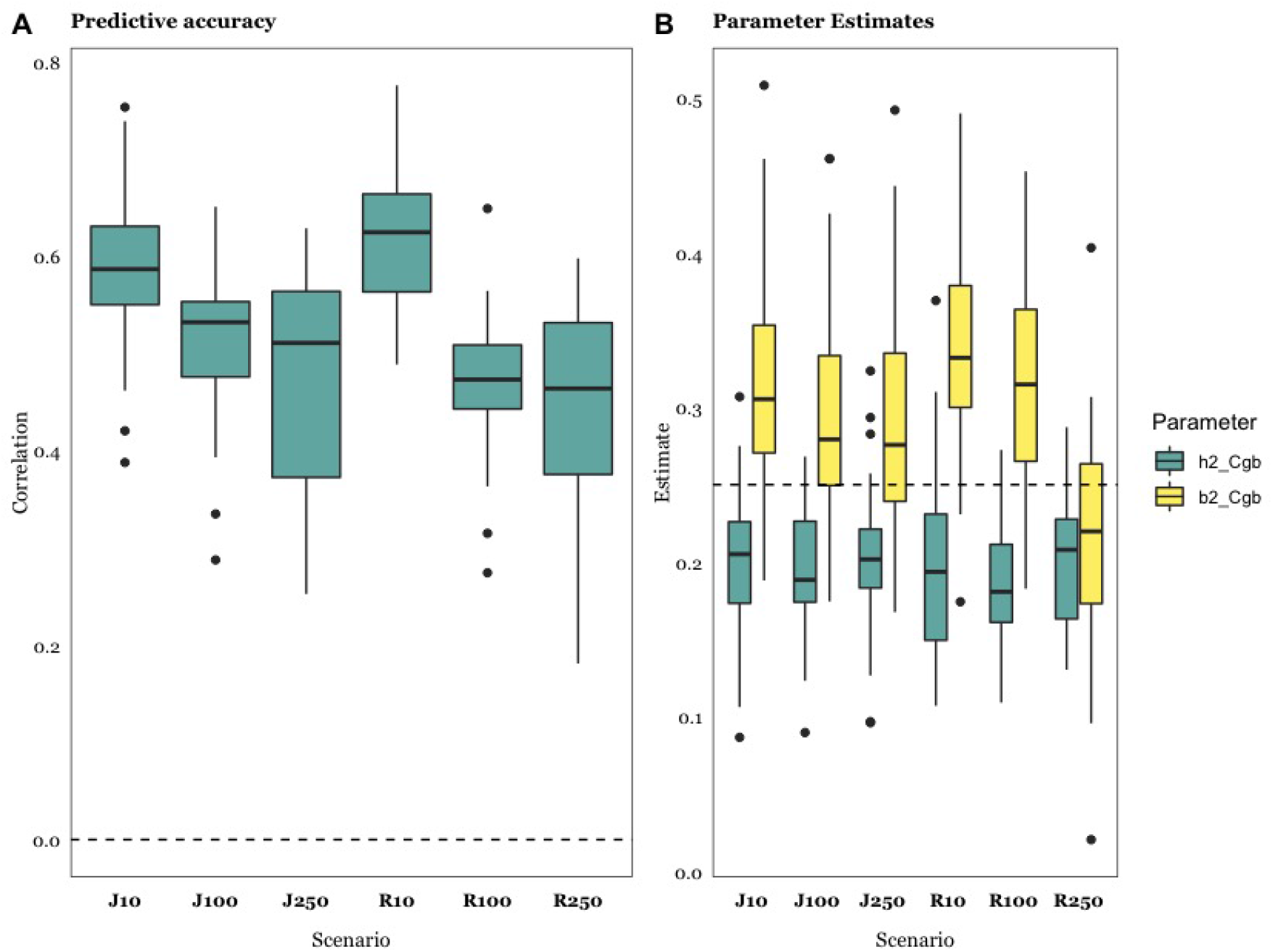
Effect of varying number of causative OTUs, r^2^ = 0.5. **A**: Predictive accuracy, computed as correlation between predicted and observed phenotypes, using Bayes Cgb. **B:** Heritability and microbiability estimates using Bayes Cgb. Details of scenarios are in Table 2 and diagrams in Fig 1: Jx, Joint scenario; Rx, Recursive scenario, with x being the number of causative OTUs (x = 10, 100, 250), half of them under genetic control. Results shown are the average of 30 replicates.

**Table 2.** Scenarios used to evaluate sensitivity to the number of causative OTUs. Symbols as in Table 1.

| Scenario | Abbreviation | $N_{\text{QTN}}$ | $N_{\text{OTU}}$ | $N_{\text{OTU(g)}}$ | $r^2$ | $h^2$ | $b^2$ |
| --- | --- | --- | --- | --- | --- | --- | --- |
| Joint | J10 | 100 | 10 | 0 | 0.50 | 0.25 | 0.25 |
|  | J100 | 100 | 100 | 0 | 0.50 | 0.25 | 0.25 |
|  | J250 | 100 | 250 | 0 | 0.50 | 0.25 | 0.25 |
| Recursive | R10 | 100 | 10 | 5 | 0.50 | 0.25 | 0.25 |
|  | R100 | 100 | 100 | 50 | 0.50 | 0.25 | 0.25 |
|  | R250 | 100 | 250 | 125 | 0.50 | 0.25 | 0.25 |

In all, our results suggest that predictive accuracy could be increased by ~ 50 % when considering microbiome data, provided microbiability is of the same order as heritability (Fig 3). We speculate that this is probably an upper limit, since it will be difficult to have microbiome data collected homogeneously across time and in different locations. While individuals can be genotyped at birth, the microbiome in early life is not representative of adult or later stages. Maltecca et al., for instance, show that early life microbiota is not a good proxy for carcass composition in pigs [35].

We observed, roughly, a two-fold increase in predictive accuracy when doubling heritability for Genome, Joint and Recursive scenarios, and a 50% increase for Microbiome or Indirect scenarios (Fig 3A vs. 3B).

### Are microbiability estimates reliable?

Reliable parameter estimates are needed to optimize the design of breeding schemes or microbiome wide association studies (MWAS) [36]. They are also needed for understanding the biology behind the interaction of microbiome and complex phenotypes. Thus far, microbiability has been usually estimated using ‘standard’ linear methods, e.g., [4,9,27], much as we have done here. It is of interest then to know how accurate these estimates could be.

Fig 4 shows estimates of variance components for each of the scenarios in Table 1. *Bayes Cgb* allows us to assess whether *h*^*2*^ and/or *b*^*2*^ are different from zero: microbiability estimate is near zero when the data are simulated according to the Genome scenario and heritability is zero when the Indirect or Microbiome scenarios hold, as it should. Similarly, both *h*^*2*^ and *b*^*2*^ estimates are near zero when the null scenario holds (Fig S1B). An overestimation of *b*^*2*^ is nevertheless evident in Fig 4, and it does not vanish at higher *r*^*2*^. This upward bias in *b*^*2*^ estimate is accompanied by an underestimation of *h*^*2*^, indicating that variance estimates are confounded when using *Bayes Cgb* model. This bias decreases though when the number of causative OTUs increases. For instance, the bias in *b*^*2*^ estimate is ~ 40% when N_OTU_ = 10 but is reduced to ~10% with N_OTU_ = 250 (Fig 4B). Therefore, it is likely that the presence of a few causative OTUs, but of large effect, combined with the presence of highly leptokurtic abundance distributions, may result in biased parameter estimates. This should be considered when interpreting microbiability estimates in real experiments. For instance, Difford et al. [4] report estimates *h*^*2*^ = 0.21 and *b*^*2*^ = 0.13 (N = 750), finding **G** and **B** to behave independently. Assuming the number of causative OTUs is small compared to that of SNPs with an effect on abundances (QTNs), we can presume Difford’s estimate of *b*^*2*^ to be inflated. This means that the actual microbiome contribution may be too small to improve prediction over that obtained from using marker data exclusively. Although authors focused on inference and not so much in prediction, Difford et al reported that no bacteria genera were significantly associated with methane emissions [4]. Other authors in turn have reported polymicrobial associations, including members of bacterial, archaeal, fungal, and protozoan communities, with methane emissions, e.g., [11,25,37–39].

For comparison, Fig 5B,D show the estimates obtained with *Bayes Cg*, when only *h*^*2*^ is estimated, or *Bayes Cb*, only *b*^*2*^ is estimated. The most noticeable aspect is that bias in *b*^*2*^ estimates is somewhat reduced relative to that found with *Bayes Cgb*, signaling again some confounding between *b*^*2*^ and *h*^*2*^. Bias was reduced overall at higher *r*^*2*^ but did not vanish.

**Fig 5:**
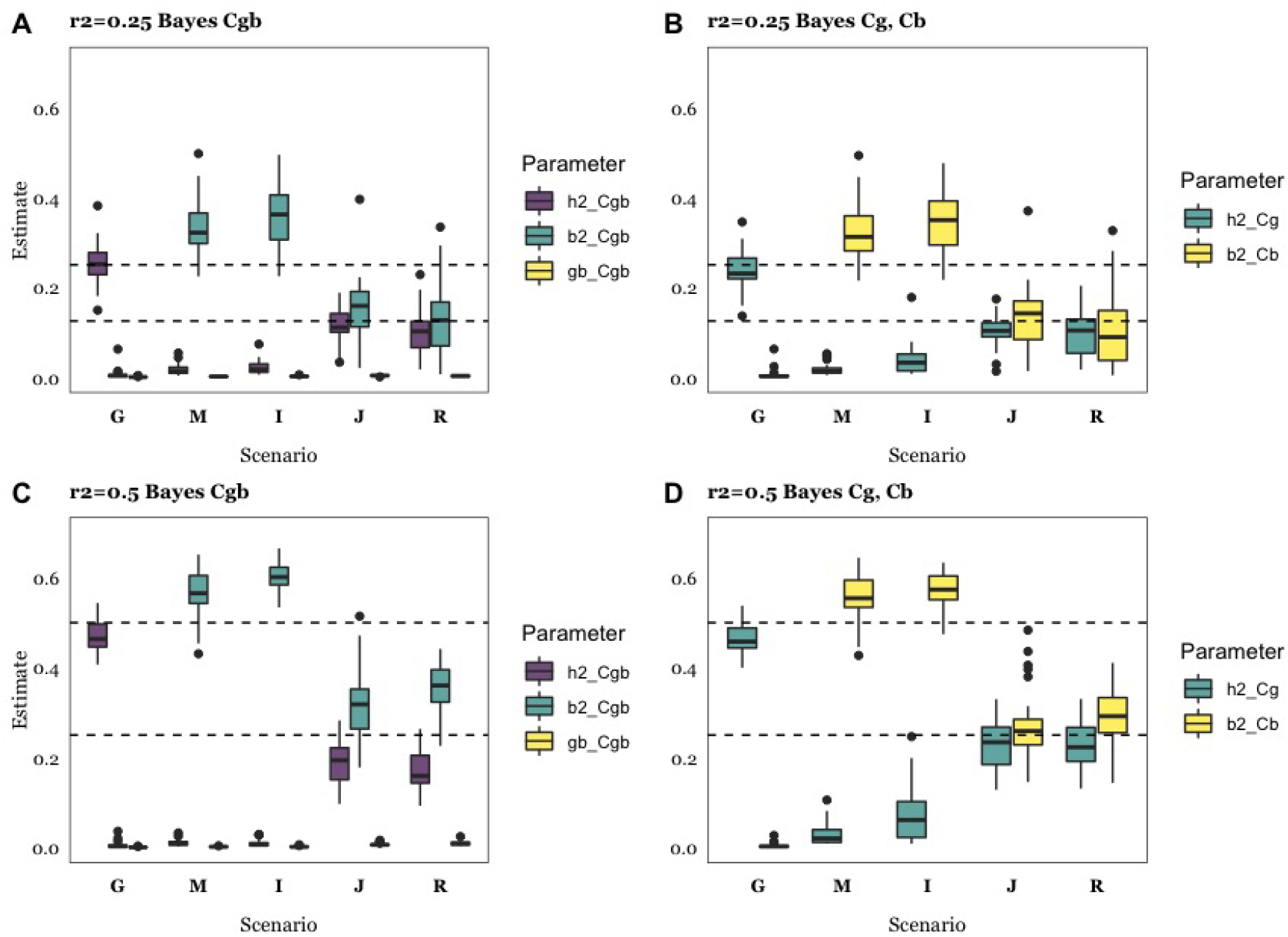
Estimates of heritability (h2), microbiability (b2), and correlation between genome and microbiome (gb) for each of the three Bayes C analysis models: Cgb includes microbiome and genome in the model (left panels); Cg includes genome only, and Cb includes microbiome data only (right panels). Upper rows correspond to r^2^ = 0.25 and lower rows to r^2^ = 0.50. Details of simulation scenarios are in Table 1: G, Genome; M, Microbiome; I, Indirect; J, Joint; R, Recursive. Horizontal dashed lines indicate true h^2^ or b^2^ parameter values (0.125, 0.25, 0.5 depending on the scenario and on r^2^). Results are average of 30 replicates. **A**: r^2^=0.25, Bayes Cgb estimates (h^2^, b^2^ and gb); **B**: r^2^ = 0.25, Bayes Cg (h^2^) and Cb (b^2^) estimates; **C**: r^2^ = 0.50, Bayes Cgb estimates; **D**: r^2^ = 0.50, Bayes Cg and Cb estimates. Data are average of 30 replicates per case.

### Can the underlying biological scenario be recovered? Can causative OTUs be identified?

An important goal of many experiments is to dissect the biological basis of microbiome and genome interactions, even if this is not strictly needed for prediction. So far, our simulations suggest that standard statistical methods can be used to quantify – with some bias – microbiability contribution to phenotypic variance. It also seems feasible to distinguish whether the Microbiome or Genome scenario fits real data best. Similarly, it seems plausible to assess when **G** and **B** contribute to the phenotypic variance, i.e. when Recursive or Joint scenarios are plausible.

Could Joint vs. Recursive scenarios be distinguished? Can data point to which of Indirect or Microbiome scenarios is more plausible, if any? Further, can causative OTUs be identified? These are far more difficult questions to answer than assessing prediction performance or estimating microbiability. Compare variance component estimates obtained under the Joint or Recursive scenarios (Fig 5): they are nearly identical for the same *r*^*2*^. The two scenarios differ in that at least some causative OTUs abundances can be under partial genetic control in the Recursive scenario. The Recursive scenario should result in a covariance between **G** and **B**. We conjectured that the two scenarios could be distinguished by analyzing the covariance*Cov*(***u***^*(i)*^, ***v***^*(i)*^) / *Var(**y**)* (see methods). Unfortunately, these estimates are close to zero irrespective of the true scenario (Fig 5A, C). The likely reason is that the actual fraction of phenotypic variance explained by indirect effects is *conditionally* negligible. Note there can be a genetic effect of **G** on **B** but, for our purposes, we are interested only in those genes that affect causative OTUs (i.e., those that affect the phenotype) and not on the whole microbial system.

An alternative approach to infer whether the Recursive causative scenario holds or not is to run a genome-wide association study (GWAS) for each of the OTU abundances on each SNP, where the SNP P-values can indicate a genetic basis for some of the abundances. If we identify significant SNPs for OTUs likely influencing **y**, we could conclude that the Recursive scenario is plausible. Unfortunately, this analysis can be doomed by the large number of tests to be realized, i.e., N_OTU_ x N_SNP_. To illustrate the caveats of GWAS on abundances, Fig 6A shows the distribution of −log10 P-values of neutral SNPs vs. SNPs with an effect on abundances. Assume we take the 5% empirical threshold of the neutral P-value distribution as indicative of association. Simulations suggest that only ~3% of causative SNP P-values will be above that threshold, i.e., approximately what is expected by chance. These P-values depend of course on the actual number of causative SNPs and on abundance heritabilities, but most evidence so far points to a weak relationship between genome and microbiome [22]. We warn it is going to be very difficult to identify abundance causative SNPs using GWAS information alone [9,20].

**Fig 6:**
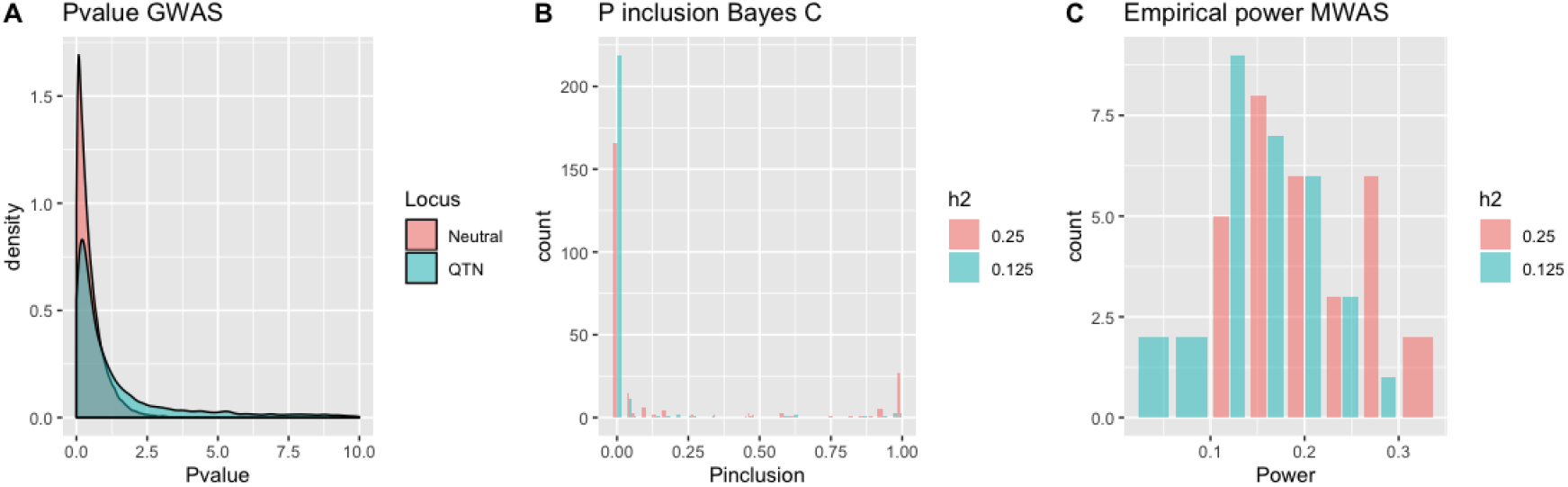
**A)** Distribution of −log10 P-values of a GWAS of abundances on SNP data; **B)** Probability of inclusion in the Bayes Cgb model of causative OTUs for the two levels of microbiability considered. **C)** Power of identifying a causative OTU computed as the probability of exceeding the 95% threshold of the empirical distribution of P-values in an MWAS for the Recursive scenario.

Another question of interest is how many of the OTUs affecting the phenotype can we expect to discover. One option is to count the frequency of a given OTU entering into the Bayes C model during sampling. Fig 6B shows the probability of including a causative OTU in the Bayes C sampling chain, which varied between ~5% (*b*^*2*^ = 0.125) to ~20% (*b*^*2*^ = 0.25). About 50% (*b*^*2*^ = 0.25) or 30% (*b*^*2*^ = 0.125) of causative OTUs were among the 5% most frequently included OTUs in the Bayes C chain, on average. Since the number of causative OTUs was 25, the rate of false positives was high nevertheless. We can conjecture that only a few causative OTUs are likely to be identified in medium-sized experiments, such as this one.

An alternative approach is a Microbiome Wide Association Study (MWAS), i.e., to perform a linear regression of the phenotype on each of OTU abundances and then select the significant results as potential causative OTUs [4]. Fig 6C shows the average power, defined as the percentage of true causative OTUs within the 5% most significant results. Power was ~15% and ~20% for *b*^*2*^ = 0.125 and 0.25, respectively, in the Recursive scenario. Again, this is not too satisfactory, as we expect a high fraction of false positives. In this particular scenario, it is perhaps more useful to consider probabilities of inclusion in the Bayes C chain rather than at P-values since the former are the result of a joint analysis of all OTUs and can be used directly for prediction.

Finally, we investigated the pattern of abundance heritabilities. Fig 7A shows the simulated heritabilities for the causative, inherited OTUs, which approximately follows a gamma distribution, together with estimated heritabilities for the causative OTUs in the Recursive scenario. We observe that both distributions are rather similar although estimates are somewhat shrunk towards zero, a consequence of using a REML-like prior. A problem of course is that we do not know which OTUs are inherited and which are not, and the true distribution of OTU heritability estimates will be a mixture. Fig 7B illustrates the heritability distributions of neutral (non-inherited) and causative (inherited) OTUs. In Fig 7B, we mixed 1.7 neutral OTU per causative OTU. This is completely arbitrary since we do not know the actual number of OTUs under genetic control, but we did so because the resulting mixture is similar to the distribution of heritabilities observed by Difford et al. (Fig 7C). If distributions in Fig 7B were representative of the true state of nature, this would suggest that about 1/(1+1.7) ~ 40% rumen OTUs could show some genetic additive variance in the experiment reported by Difford et al.[4].

**Fig 7:**
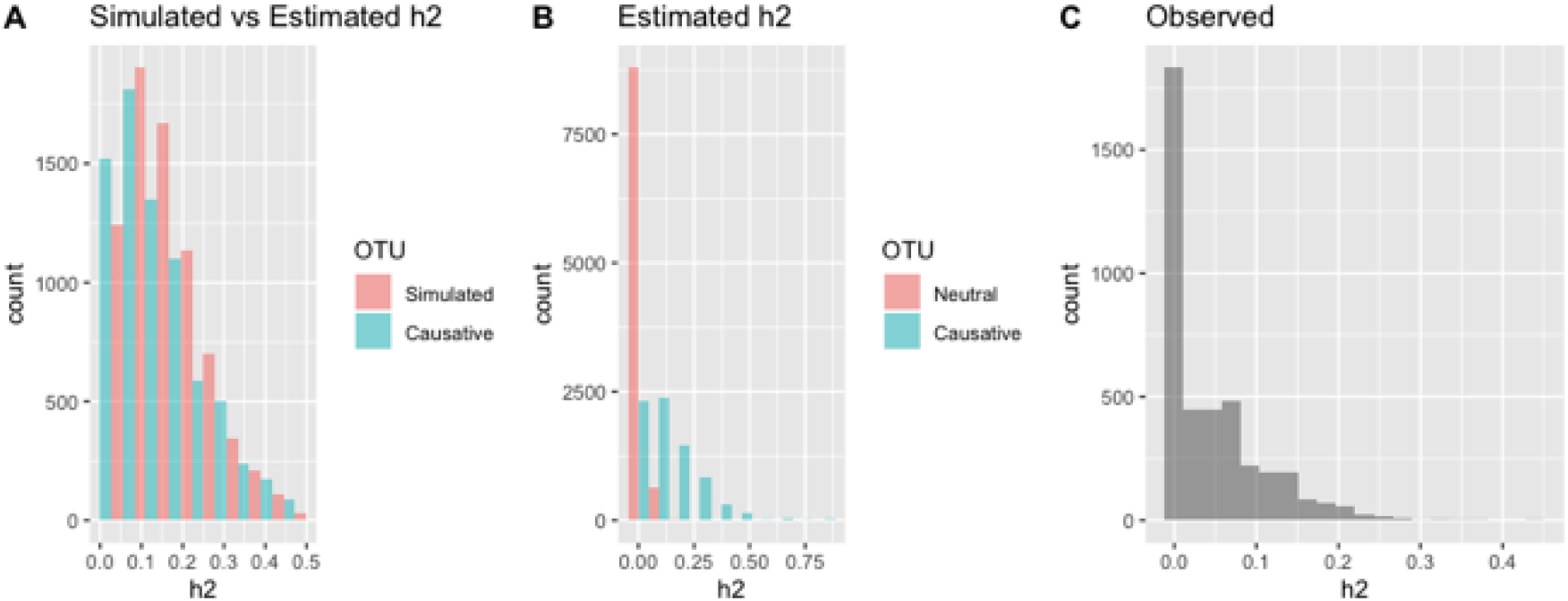
**A:** ‘True’ (simulated) and GBLUP estimated distribution of abundance heritabilities for causative OTUs in the Recursive scenario. **B:** GBLUP estimated distribution of abundance heritabilities for neutral and causative OTUs in the Recursive scenario. **C:** Actual distribution of OTU abundance heritabilities reported by Difford et al.[4].

## Discussion

Fig 1 represents but highly simplified relationships between the genome, microbiome, and phenotype. These scenarios are nevertheless important to interpret empirical data and can help to identify limiting factors in prediction. Further, provided a good fit is found, they will help in designing experiments that combine microbiome and genetic data. We chose parameter combinations that represent extreme case scenarios and we found that results were, qualitatively, robust to parameter choice such as *r*^*2*^. A parameter that can be relevant though is the number of causative microbiome taxa, i.e., those with an effect on the phenotype. This number seems to affect the bias of microbiability estimates (Fig 4).

Here, we have proposed a new simulation procedure that addresses some important challenges. First, the algorithm avoids the need for actual phenotype simulation by using real genotype and abundance data. Although we concede that this procedure may limit the generality of the study, e.g., in terms of data size, we believe the advantages of using real data are numerous, since no simulation procedure can accommodate all known and unknown subtleties of the highly dimensional distributions at hand. Second, we develop an ingenious permutation procedure (Box 1) that allows linking previously uncorrelated data to fit a desired genetic hypothesis. By also permuting all OTUs within a given cluster, we minimize disruption of the whole covariance structure (Fig S2).

Numerous studies have reported microbiability values for economically important traits e.g. [4,12,25,39], but their actual reliability is not known. Estimates may be affected by the estimation procedure. There are numerous alternatives to estimate *b*^*2*^, among them Bayes C [15], GBLUP [40], Bayesian RKHS regression using either Bray–Curtis dissimilarities as relationship matrix [25] or with the variance-covariance from the log-transformed OTUs as kinship matrix[25,41]. Our results (Fig 5) indicate that BayesC estimates may be biased upwards, especially when *b*^*2*^ is higher than 0.25 and the number of causative OTUs is small. However, we found that estimates of *b*^*2*^ derived with Bayes C were very close to zero in the null scenario (Fig S1B); therefore, we conclude that models using priors from the Spike-Slab family, which contemplate a priori the possibility of null effects, can be used to test whether heritability or microbiability is substantial. Ramayo-Caldas et al. [25] report that estimates using Bray-Curtis based kernels are higher than those using the log-transformed covariance matrix. The behavior of estimation methods for microbiability merits further research.

One conclusion from this work is that it is going to be difficult to distinguish between some underlying scenarios or to identify the causative OTUs and SNPs, at least using standard linear models as was done here. The distinction between Joint and Recursive scenarios is of special relevance for breeding. The latter assumes partial genetic control of some causative OTUs. Yet, we found both scenarios result in very similar patterns (Figs 3, 4, 5). Perhaps, a more powerful approach would be to use structural equation models (SEM), which allow including a variable both as independent and dependent. Saborio-Montero et al. [42] compared a linear bivariate (one OTU and the phenotype) model with a SEM but found few differences. One restriction of their approach is that one SEM was fitted for each abundance. A whole-genome approach seems in principle more adequate; however, modeling recursive effects in this context is both statistically and computationally challenging because of the large number of SNP-OTU combinations that would need to be considered.

A line of research that we have not considered involves possible microbiome-DNA interactions. Although the number of possible interactions to consider can be huge when the number of SNPs and the number of OTUs is large, interactions between features in two high-dimensional sets can be modeled in a Gaussian context using co-variance functions. These functions are the Hadamard product of set-specific similarity matrices such as the Hadamard product of a SNP-derived and an OTU-derived ‘relationship’ matrix. Such an approach has been used before to model, e.g. interactions between SNPs or between SNPs and environmental covariates (e.g., 43).

The usefulness of microbiome in prediction depends crucially on its stability in time and space. For instance, although measures of gastrointestinal microbiome abundances are known to be repeatable, it cannot be expected to remain stable throughout an individual’s life span. After weaning and under standard management conditions, e.g., constant diet and absence of antibiotic treatment, the diversity of monogastric gut microbiota increases with host age until its composition remains stable. Rumen microbial communities are highly resilient and host-specific [44,45] but change in early life. The transition towards a more stable an adult-like ruminal ecosystem occurs between weaning and one year of age [46]. Therefore, for prediction purposes, we recommend the inclusion of microbial information at least after weaning, preferably at adulthood. This may limit the usefulness of microbiota for prediction in breeding schemes as compared to genomic data in livestock.

At present, modeling the influence of microbiome abundances on complex phenotypes is an open area of research. Here we have presumed that the effects on abundances are additive in the log scale. Similar models are widely used in a diversity of scenarios, e.g., multiplicative models are used to accommodate fitness effects in evolutionary genetics [47] or to deal with highly leptokurtic distributions such as raw abundances, an effect that is smoothed with the log transformation. In addition to the log-transformation, a widely popular choice in genetics is the threshold model [7], which assumes the presence of a continuous liability (here abundances) with an effect value ‘0’ below a given threshold and ‘1’ otherwise. This model has the advantage of being independent on whether abundances are log-transformed or not and is also biologically sound since it is conceivable that a minimum microorganism abundance is required to trigger a particular effect. To test the robustness of the log-transformation, we simulated phenotypes such that 25% of causative abundance observations were above the threshold and the analysis was performed on the log transformed abundances as before. As could be expected, using a ‘wrong’ model for the analyses was detrimental to prediction but not dramatically (Fig 8A). Parameter estimates were affected downwards compared to the multiplicative model (Fig 8B). We suggest that major conclusions from this work should hold even if the relationship between variables and phenotype is not strictly multiplicative.

**Fig 8.**
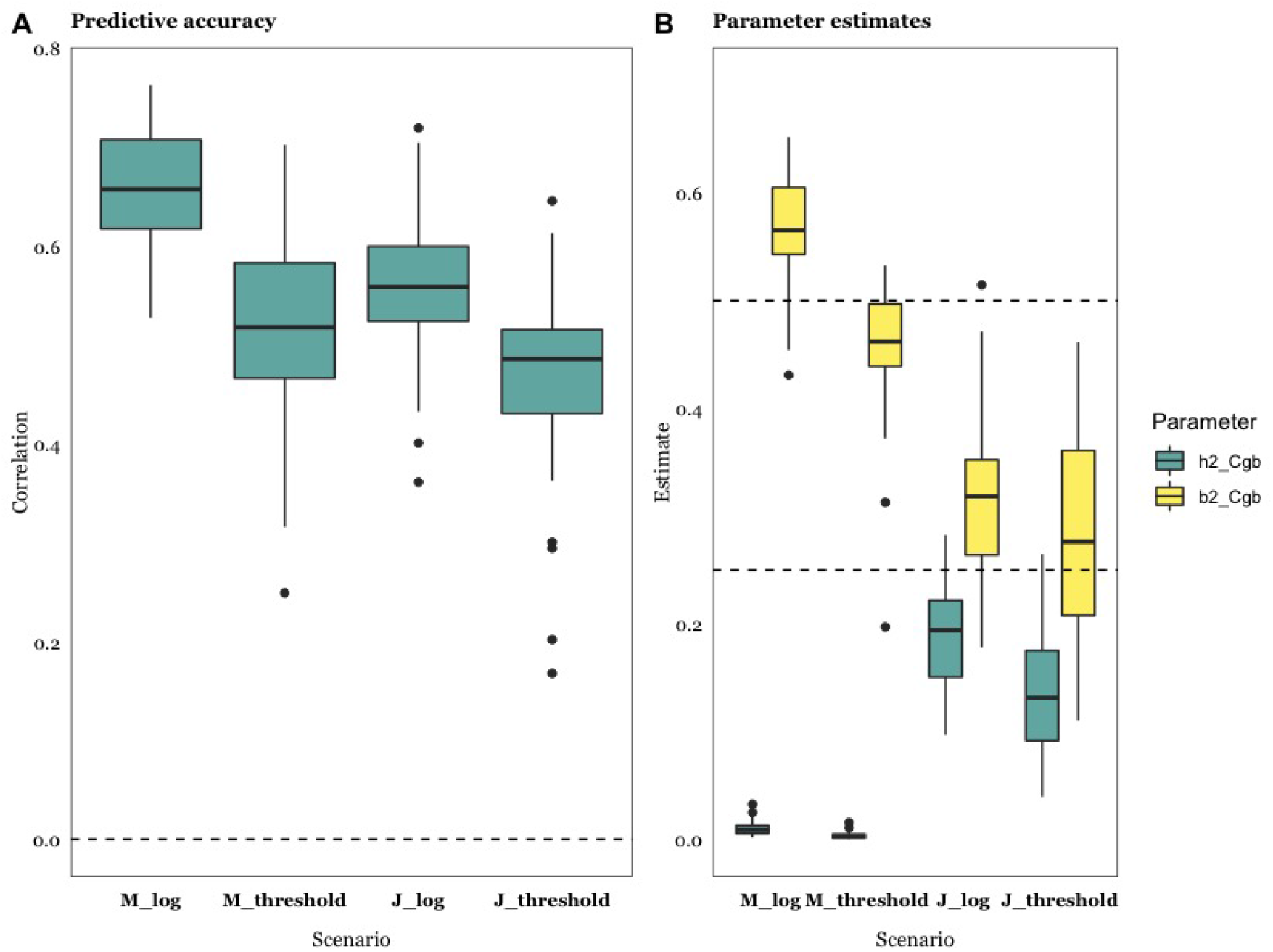
Comparison of multiplicative (log) and threshold Microbiome (M) and Joint (J) scenarios (r^2^ = 0.5). **A**: Predictive accuracy, computed as correlation between predicted and observed phenotypes, using Bayes Cgb. **B:** Heritability (h^2^) and microbiability (b^2^) estimates using Bayes Cgb. Results are average of 30 replicates. Scenarios M and J as specified in Table 1; the log transformation results are shown for completeness and are the same as in Figs 3 and 5. Data are average of 30 replicates.

## Conclusion

This study suggests that microbiome data can significantly improve the prediction of complex phenotypes, irrespective of whether some abundances are under direct genetic control or not. For this strategy to be successful, though, medium to large-sized experiments are required, the microbiome should be relatively stable and should be available before the phenotype is collected. This limits the usefulness of microbiome for prediction in breeding schemes as compared to genome data, which can be collected at birth and remains unchanged. Important potential applications remain nevertheless, such as predicting methane emission in cattle, feed efficiency, disease predisposition, or crop production using soil metagenome. Overall, we can be rather confident that standard linear methods can be used despite the highly leptokurtic distributions observed in OTU abundances. There is room for specific theoretical developments though, perhaps along the lines proposed by Saborio-Montero et al. [42], but these should be based on a better understanding of the relation between microbiome and phenotype. It seems critical to quantify, even approximately, the number of taxa affecting the phenotype and to characterize the distribution of their effects. We are far less optimistic in what regards the identification of causative OTUs, and in particular of the putative QTNs affecting relative abundances.

## Materials and Methods

### Simulation Strategy

There is ample literature and software available on the simulation of ‘standard’ complex phenotypes, e.g., [48–51]. These algorithms, however, are not suited for some of the scenarios posed in Fig 1. Here we propose simulating the joint influence of genome and microbiome on a quantitative trait by adding their contributions plus a random noise:

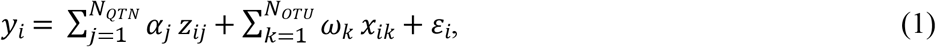

where *y*_*i*_ is the i-th individual record, *α*_*j*_ is the genetic effect of j-th causal SNP (QTN), with *j* = 1, N_QTN_, the number of QTNs, *z*_*ij*_ is the genotype of the i-th individual for j-th SNP coded say −1, 0 and 1 (strict additivity was assumed for all QTN), *ω*_*k*_ is the linear effect of the k-th OTU abundance (*x*_*ik*_), with *k* = 1, N_OTU_, the number of abundances that influence the phenotype and ε is a normally distributed residual. The OTU’s coefficient can be interpreted as the expected change in phenotype per OTU’s abundance unit increase. Since abundances are in the log scale, this is equivalent to a multiplicative effect model. Equation (1) is valid for all scenarios in Fig 1, except that the term involving markers 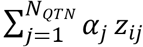 is removed in the Microbiome and Indirect scenarios whereas the term 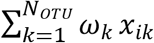 is removed in the Genome scenario.

For the Indirect and Recursive scenarios, we also need to model the variation in abundances (*x*) that is explained by the genome (Fig 1). Again, we can resort to a linear model where the abundance itself is treated as a standard complex phenotype:

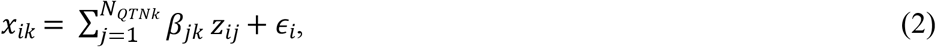

where *x*_*ik*_ is the abundance level of the k-th OTU that is under partial genetic control for i-th individual, *β_j_* is the genetic effect of j-th QTN on abundance, and *z*_*ij*_ is the genotype of the i-th individual for j-th SNP. The j-th sum is across the QTNs influencing k-th abundance, j = 1, N_QTN(k)_. Note abundances *x*_*ik*_ in Eqn. (2) are a subset of those in (1). There may be other non-causative abundances under genetic control, but this is irrelevant for our purposes. A phenotype following the Recursive scenario can then be simulated via a two-step procedure: first, simulate abundances (*x*) using Eqn. (2) followed by phenotype simulation using (1) given the abundances obtained.

We used real genome and microbiome data as input for the simulation procedure. We downloaded the rumen abundance table of 4,018 OTUs from dairy cattle rumen (N = 750, [4]). A pseudo-count equal to one was added to zero abundances, which were next total-sum scaled and log-transformed. This results in much less leptokurtic and less asymmetric distributions than original raw abundances. In Eqns. 1 and 2, *x*_*ik*_ represent the already log-transformed abundances. As for genotypes, high-density array genotypes from 750 dairy cows among the total available were downloaded from [11]. To prune SNPs and facilitate computation, 35% of all genotypes with a minimum allele frequency of 0.01 and a maximum missing percentage of 1% were retained. A total of 32,204 autosomal SNPs was finally retained. The few missing values were simply imputed with the mean. Thirty simulation replicates per scenario were simulated.

Under the Joint scenario, which assumes independence between **G** and **B**, we can simply sample the list of causative SNPs and abundances, simulate their effects, and apply Eqn. 1 to generate phenotype values given observed genotypes and abundances. The case of Recursive and Indirect scenarios is not that obvious because we need to sample abundances that are under genetic control and a link must exist between **G** and **B** (Eqn. 2). We solved this issue by rearranging abundances of a given OTU between individuals such that the desired correlation between abundance and individual’s genotypes is attained. This strategy has the important advantage that the distribution of abundances is not changed. Suppose 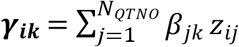 is the simulated genetic effect of the i-th individual for k-th abundance (Eqn. 2) and that the desired heritability for that abundance is 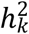. The algorithm (Box 1) is based on the simple observation that, given any two vectors *x* and *y*, correlation is maximum (*p* ~ 1) when observations in both vectors are sorted and *p* is ~ zero when they are shuffled. Therefore, there must be some order ***y*_*sort*_** then that fulfills, approximately, the constraint cor(***x*, *y*_*sort*_**) = *p*. For our purposes, we need to rearrange the observed abundances ***x*_*k*_** such that the correlation between rearranged ***x*_*k*_** and ***γ*_*k*_** is *h*_*k*_, the square root of heritability for *k*-th abundance. The algorithm is detailed in the Box.

A drawback of this algorithm is that it locally breaks the covariance between abundances of different OTUs. To alleviate this, we permuted all abundances that fell within the same OTU cluster. We clustered abundances using R function hclust(dist(.), method=“ward.D2”) and cut the tree in K = 500 clusters. We chose K = 500 because the first quartile of intra-cluster average correlation was above the third quartile of the average correlation between random abundances, that is, clusters were made up of highly correlated abundances compared to average. We also explored K = 200 but we did not find any difference neither in predictive accuracy nor in heritability estimates. To verify that the shuffling algorithm did not alter the whole structure of the data, we show the principal component analysis of the original and a few shuffled microbiome sets in Fig S3.

**Algorithm 1:**
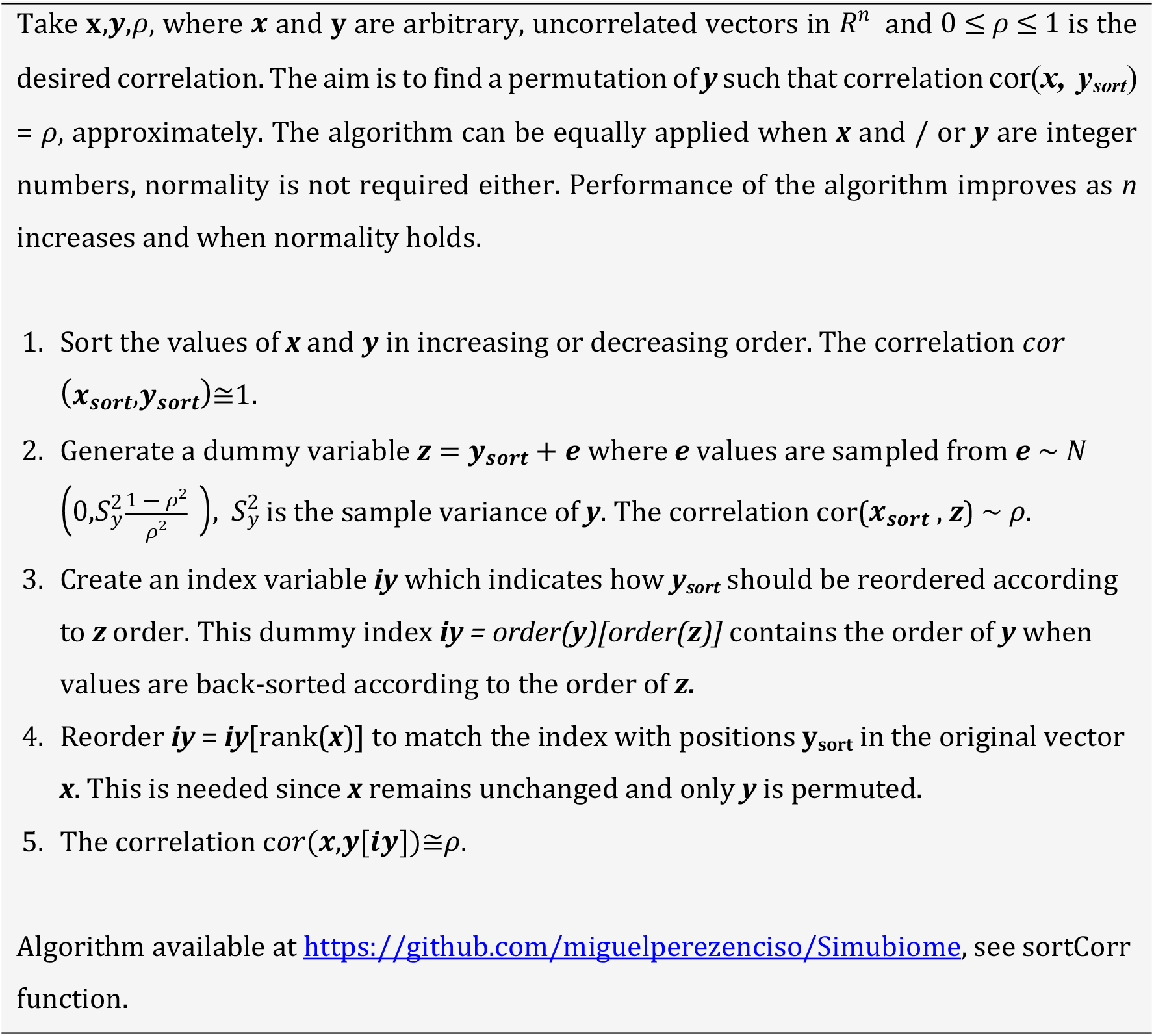
Find a permutation of vectors *x* and *y* such that the correlation between permuted vectors is a predetermined value ρ.

### Parameter fitting

Little is known neither on the number of OTUs influencing a given phenotype nor on how many of those are partly inherited. For that reason, we chose some extreme, yet ‘educated’ values for each of the five scenarios depicted in Fig 1. We considered 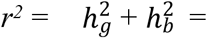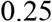 and 0.50; *r*^*2*^ = 0.25 is grossly the value reported by Difford et al. 2019 with N = 750, whereas values closer to *r*^2^ = 0.50 were reported by Wallace et al. in some farms. Overall, augmenting *r2* values tries to mimic the effect of increasing sample size. We assumed 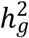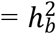 for Joint and Recursive scenarios, as also reported by Difford et al or Camarinha-Silva et al. approximately. The number of QTNs was fixed to 100. This figure is somewhat arbitrary, but the specific number of loci would not affect much the results. Barton et al. [52] showed theoretically that most properties of the infinitesimal model converge as fast as the inverse of the number of loci, or ~ 1% deviance with N_QTN_ = 100. In general, genomic prediction is known to be relatively insensitive to the number of QTNs [53]. As for individual genetic effects α, numerous empirical and theoretical works show that they are not uniformly distributed and can be approximated by a gamma-like distribution [54,55]. Here we sampled genetic effects α ~ Γ(shape = 0.2, scale = 5), as suggested by Caballero et al. [56], and also used previously by us [57].

Much less is known on the number of causative OTUs (N_OTU_), although we can presume that N_OTU_ should be smaller than the number of QTNs. For instance, Duvallet et al. [36] found in a large meta-analysis that the human diseases studied were affected on average by 10 - 15 changes in abundances at the genus level. Here we considered N_OTU_ = 25 (0.6% of all OTUs), although we also evaluated N_OTU_ = 10, 100 and 250. Similarly, for the Recursive and Indirect scenarios, we took the extreme scenario where all causative OTUs are genetically determined, i.e., N_OTU_ = N_OTU(g)_. The genetic effects *β* on abundances (Eqn. 2) were sampled from the same distribution β ~ Γ(shape = 0.2, scale = 5) as direct genetic effects α. We are much more ignorant regarding the distribution of abundances’ effects ω on the phenotype (Eqn. 1). We took as proxy the regression coefficients of methane emission on abundances published by Difford et al. [4], in their supplementary information S4, which can be approximated by a Γ(shape=1.4, scale=3.8). Fig S3 compares both gamma distributions and the fit to empirical data. This model predicts that the variance of OTUs’ effects is wider and of larger individual effect on average than that of SNPs. Although this is speculative at this point, it is sensible to assume that only a few taxa do have a sizeable influence on the phenotype, say methane emission.

### Analysis

We used Bayes C algorithm [15] as implemented in BGLR [58] to assess prediction performance and reliability of parameter estimates. We also tested Bayesian RKHS regression, equivalent to GBLUP [40], but results were similar or worse and are not presented. Three models were used to analyze the data:

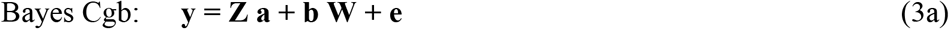

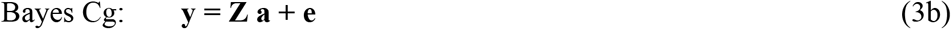

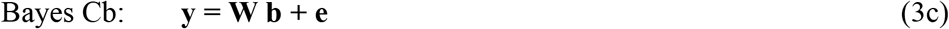

where **y** is the vector containing the simulated phenotypes, **a** contains the marker effect estimates, **Z** contains the observed genotypes for the 33k markers, **b** contains the OTU abundance effects, **W** is a matrix with all 4,018 abundances in the 750 individuals, and **e** is the residual. Prior to the analyses, phenotypes, abundances, and genotypic values were standardized to mean zero and SD = 1. As priors π for SNPs or abundances probabilities to enter into the model, we used π ~ Beta(p_0_ = 5, π_0_ = 0.001), which has expectation π_0_ and variance π_0_ (1−π_0_) / (p_0_+1). We also considered a much more liberal, flat prior for π ~ Beta(p_0_ = 2, π_0_ = 0.01), but we did not observe strong differences. A total of 50k iterations were run per Bayes C chain, a plot of the residual variances along iterations indicated convergence was attained with this number of iterations. To assess predictive accuracy, 75 (10% of N) phenotypes were randomly removed and predicted with the fitted model. Correlation between observed and predicted phenotypes was used as measure of predictive accuracy.

The ‘heritability’ is not explicitly defined in a Bayes C framework, and here we used the proposal by [58] (https://github.com/gdlc/BGLR-R/blob/master/inst/md/heritability.md). In short, at each iteration *i*, the algorithm samples SNPs and OTUs effects:

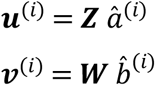

where ***u***^(*i*)^ and ***v***^(*i*)^ are genome and microbiome effects at i-the iteration for the set of individuals, respectively, 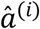 and 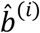 are current SNP and OTU abundances solutions; therefore, *Var*(***u***^(*i*)^) / *Var*(***y***) and *Var*(***v***^(*i*)^) / *Var*(*y*) are i-th iterate heritability and microbiability estimates wherefrom posterior means can be estimated by averaging over iterations. For Bayes Cgb, we also sampled the absolute covariance between **u** and ***v***, i.e., |*cov*(***u***^(*i*)^, ***v***^(*i*)^)| / *Var*(***y***).

To assess how likely is to identify causative OTUs, we retained the probability of a given OTU entering into the model, averaged over Gibbs sampling iterations. We run a GWAS of abundances (*x*_*k*_, k=1, N_OTU_) on SNP genotypes (*z*_*j*_, j=1, N_SNP_) using R function lm(*x*_*k*_ ~ *z*_*j*_) and we computed the P-value of both causative QTNs, i.e., affecting abundances, and neutral SNPs. This was done in the Recursive scenario only. In this scenario, we also computed the heritabilities of all abundance levels using GBLUP via a RKHS strategy (https://github.com/gdlc/BGLR-R/blob/master/inst/md/GBLUP.md#RKHS) using BGLR. Weakly informative priors for variances were used to mimic a REML-like estimator.

## Author contributions

MPE, GDLC and LMZ conceived research. MPE and LMZ performed research. All authors discussed research. MPE wrote the manuscript with help from the rest of authors.

## Acknowledgments

This work was developed while MPE and LMZ spent a research stay at Michigan State University kindly funded by GDLC. LMZ is supported by a Ph.D. grant from the Ministry of Economy and Science (MINECO, Spain), MPE is funded by MINECO grants AGL2016-78709-R and PID2019-108829RB-I00, from the EU through BFU2016-77236-P (MINECO/AEI/FEDER, EU) and “Centro de Excelencia Severo Ochoa 2016-2019” award SEV-2015-0533. YRC was funded by Marie Skłodowska-Curie grant (P-Sphere) agreement No 6655919 (EU).

